# Effect of reversible osmotic stress on live cell plasma membranes, probed via Laurdan general polarization measurements

**DOI:** 10.1101/2021.10.21.465302

**Authors:** Elmer Zapata-Mercado, Kalina Hristova

## Abstract

Here we seek to gain insight into changes in the plasma membrane of live cells upon the application of osmotic stress using Laurdan, a fluorescent probe that reports on membrane organization, hydration, and dynamics. It is known that the application of osmotic stress to lipid vesicles causes a decrease in Laurdan’s generalized polarization (GP), which has been interpreted as an indication of membrane stretching. In cells, we see the opposite effects, as GP increases when the osmolarity of the solution is decreased. This increase in GP is associated with the presence of caveolae, which are known to disassemble and flatten in response to osmotic stress, in a process that supplies extra plasma membrane in physiological processes.

**Significance:** Cells can experience multiple stresses *in vivo*. Furthermore, the application of osmotic stress is used as a biophysical tool to interrogate membrane proocesses *in vitro*. We sought to investigate the consequences of osmotic stress on the plasma membrane properties using the fluorescent probe Laurdan. Unexpectedly, we find that osmotic stress leads to an increase in GP in live cells. The opposite change in GP has been observed in model lipid bilayers, reminding us there are limitations to the utility of model systems in understanding cell membrane behavior. Despite years of research, the cell membrane still has ways to surprise us.

## Introduction

The plasma membrane consists of a phospholipid bilayer with embedded proteins that mediate the communication with the external environment and control the transportation of nutrients and ions. The lipid bilayer is the structural backbone of the cellular membrane. For many years, it was believed to just provide a passive physical barrier to non-specific, non-protein mediated passage of molecules (1). However, recent research has revealed that the lipid components can play an active role in cell physiology (2,3). It is now appreciated that the phospholipid bilayer is highly complex and dynamic, and plays regulatory roles in many physiological processes such as cell signaling, synaptic neurotransmission, and metabolism.

Remarkably, cells possess two to three times the membrane needed to sustain their shape (4,5). This extra membrane is primarily stored in the caveolae, 60-80 nm cup-shaped invaginations (6,7). The caveolae were first observed in electron microscopy experiments as pits with flask-shaped morphology and uniform curvature. These structures play an essential role in the cellular response to acute mechanical stress as they disassemble and flatten in a rapid process independent of ATP or actin (8). Thus, caveolae serve as a membrane reservoir to protect the cell (8). This role is of great physiological significance since cells can experience multiple stresses *in vivo*. For example, epithelial cells in our lungs are exposed to cycles of mechanical stress when we breathe, endothelial cells are constantly exposed to shear flow, and myocytes endure stress and strain regularly when muscles contract and relax (9-12).

Here we investigate if the physical-chemical properties of the plasma membrane are altered upon the application of osmotic stress. We monitor changes in the membrane environment using Laurdan, a fluorescent probe that has been used for years to study lipid organization and dynamics in synthetic and biological membranes (13-39). Laurdan is an amphipathic fluorophore that incorporates into lipid bilayers and biological membranes. It is polarity-sensitive and has been used previously to monitor changes in membrane bending, tension, and/or domain formation. When excited, Laurdan undergoes charge separation, creating a strong dipole. The coupling between the Laurdan dipole and the dipoles of nearby water molecules in the membrane results in a redshift in the Laurdan emission spectrum. This hydration-induced redshift can be assessed by quantifying a single parameter, the so-called Generalized Polarization (GP) (27,35,37). The value of GP decreases as Laurdan is exposed to an increasingly polar environment and increases as the environment becomes less polar. Thus, Laurdan has been used to study the hydration of membranes. However, recent work has expanded the understanding of the Laurdan fluorescence response (40-43). It has been pointed out that free water molecules are sparse in lipid bilayers. Instead, the water molecules are tightly bound to lipid carbonyls and thus the mobility of the lipids, along with the bound water, can also affect the fluorescence properties of Laurdan. Specifically, the GP of Laurdan has been shown to increase with decreased lipid mobility and to decrease as the lipid mobility is increased (40-43).

Laurdan fluorescence has been used previously in the context of lipid vesicles that have been subjected to osmotic stress (44). In particular, a decrease in GP has been reported as a function of osmotic stress (44). These observations were interpreted as a proof for increased membrane tension and membrane stretching, which allows water molecules to enter the hydrophobic interior of the membrane. In this work, we sought to determine if similar effects are observed in the plasma membranes of live cells in response to osmotic stress.

## Experimental Methods

### Cell Culture

Experiments were performed in Chinese Hamster Ovary (CHO) and Mouse Embryonic Fibroblast (MEF) cells. The MEFs included a (+/+) version that is homozygous for the caveolin-1 gene (or wild-type) and a (-/-) knock-out.

The cells were cultured in Dulbecco’s modified Eagle medium (DMEM), supplemented with 10% fetal bovine serum (ThermoFisher), 1.8 g/L d-glucose, 1.5g/L sodium bicarbonate, and 1mM non-essential amino acids (NEEA) for the CHO cells media. Cells were seeded in 35-mm glass-bottom collagen-coated dishes (MakTek’s Corporation) at a density of 2.0×10^4. Cells were kept in an incubator at 37°C with 5% carbon dioxide.

To apply the osmotic stress, cells were incubated with a hypotonic solution (swelling media) buffered with 25mM HEPES, prepared with serum-free DMEM media (starvation media), and increasing amounts of diH2O. These different solutions contained 0, 10, 25, 40, 45, 50, 60, 65, 75, 90, 95, and 100% diH_2_O. Twenty-four hours after seeding, Laurdan from a methanol stock was pre-mixed with 1mL of starvation media to a concentration of 10μM. The cells were incubated with the Laurdan solution for 15 mins at 37°C and then washed with 1x PBS to remove the excess Laurdan. The swelling media was added to the cells at the pre-set imaging temperature at least 5mins before imaging. The temperature was adjusted using a Stable Z stage (Bioptechs).

### Vesicles

CHO cells were seeded into a six-well plate at a density of 5×10^4^, as described in (45). Twenty-four hours after seeding, cells were vesiculated as previously described (46). Briefly, cells were first rinsed three times with 1x PBS supplemented with 750μM calcium and 500μM magnesium (CM-PBS). Then the cells were incubated with 10μM Laurdan pre-mixed with the vesiculation buffer for 1 hour at 37°C. This buffer consisted of CM-PBS with 25mM formaldehyde and 0.5 mM 1,4-dithiothreitol (DTT). The formaldehyde was quenched with glycine with a final concentration of 125mM. Vesicles were transferred to a four-well glass-bottom imaging slide (MakTek) and let to settle down for 1 hour prior to imaging.

### Laurdan Spectra Acquisition and Analysis

Laurdan spectra were acquired with a two-photon microscope equipped with the OptiMis spectral imaging system (Aurora Spectral Technologies)(47). Laurdan was excited at 780 nm with a femtosecond Mai-Tai laser (Spectra-Physics). The system allows the acquisition of full spectra for each pixel in the frame, and it records the emission in a stack of 200 images at different wavelengths, approximately ∼1nm apart. To extract the Laurdan spectra, the image stacks were analyzed using the software ImageJ to plot the emission spectra in the selected membrane region of interest, as shown in Figure 1. The generalized fluorescence of Laurdan was calculated according to:

**Figure 1:**
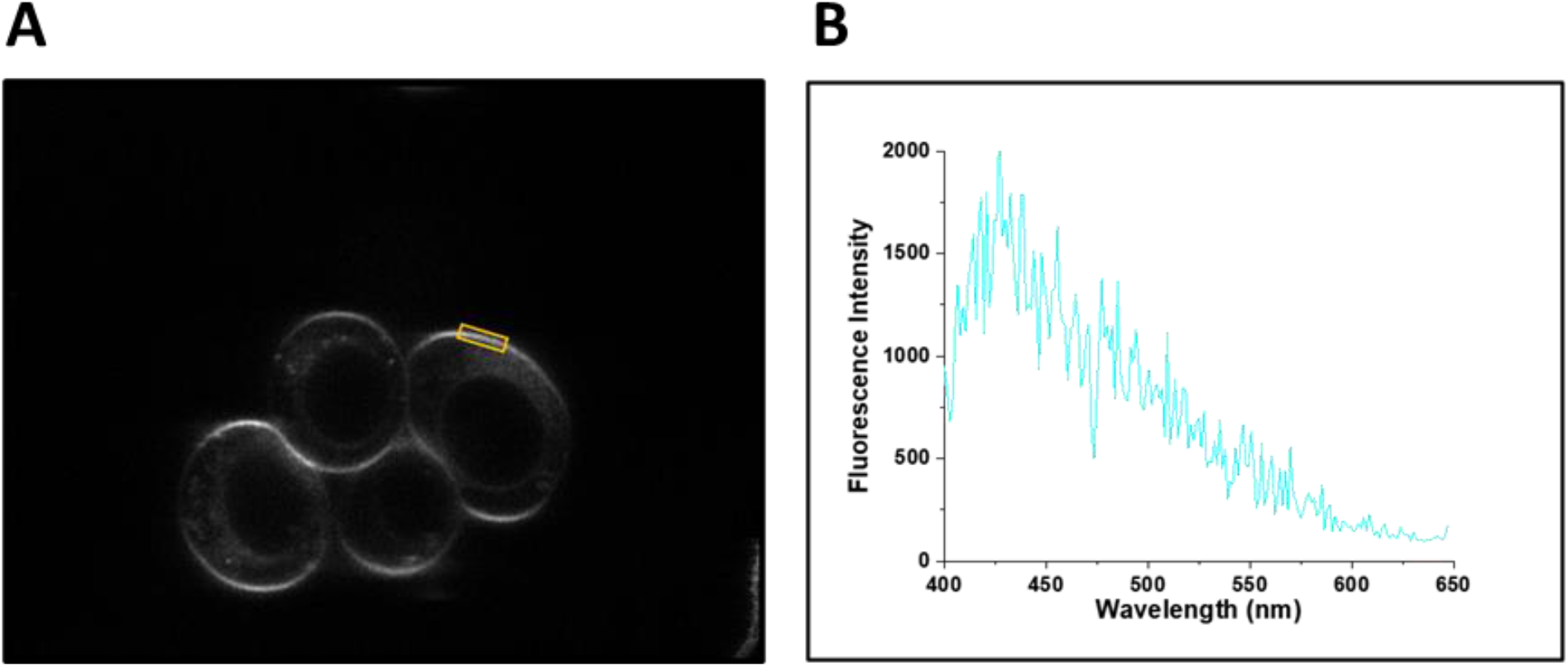
(A) CHO cells at 90% media dilution with diH_2_O. A region of interest from each cell (yellow) is analyzed and the average Laurdan spectrum is calculated and shown in **Figure 3A**. (B) Laurdan spectrum from a single pixel within the yellow region.

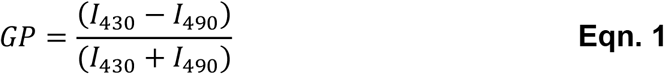

where *I*_430_ is the integrated spectral intensity in a 20nm window centered at 430nm and *I*_490_ is the integrated spectral intensity in a 20nm window centered at 490nm.

Cells (or vesicles), under all conditions studied, were imaged in at least three independent experiments. Each data set was derived from at least 20 (typically 40 to 50) individual cells or vesicles. One or two regions of plasma membrane were analyzed per cell (as shown in Figure 1A). These regions were selected randomly, from membrane areas that were in focus, and not in contact with intracellular organelles or neighboring cells.

### Statistical Analysis

The statistical analysis was based on one-way ANOVA using GraphPad Prism 8.3.

### Osmolarity measurements

Measurements were performed in the Johns Hopkins Hospital Core lab using an automated A2O osmometer (Advanced Instruments).

## Results

Here we sought to understand how the application of osmotic stress changes the plasma membrane environment. Towards this goal, we used Laurdan as a membrane probe, as its emission spectrum depends on the environment. In the experiments, 10µM Laurdan was added to CHO cells and incubated for 15 minutes. Cells were washed and subjected to reversible osmotic stress in serum-free media with increasing amounts of diH_2_O. After 5 minutes, the emission spectra of Laurdan in CHO plasma membranes were acquired as described in Methods. From these spectra, the general polarization (GP) of Laurdan was calculated using ***Equation 1***.

Integrated fluorescence images of cells at different media dilutions are shown in **Figure 2A**. The osmolarities of the media dilutions are shown in **Figure 2B**. Typical Laurdan spectra are shown in **Figure 3A**, for the cases of 0% and 90% media dilution with diH_2_O. In a standard medium (0% diH_2_O), we observe an emission maximum at ∼430 nm and a second less pronounced peak at ∼490 nm. The application of osmotic stress at 90% diH_2_O decreases the 490 nm peak. In **Figure 3B**, the Laurdan GP was calculated for all experimental conditions investigated. As diH2O in the swelling media increases, the GP stays constant up to 60% diH_2_O and increases afterward up to 95% diH_2_O. Imaging at 100% diH20 revealed a marked decrease in GP, likely indicating water penetration in the membrane. With time, many of the cells in 100% diH_2_O could not withstand the osmotic pressure, and their membranes ruptured.

**Figure 2:**
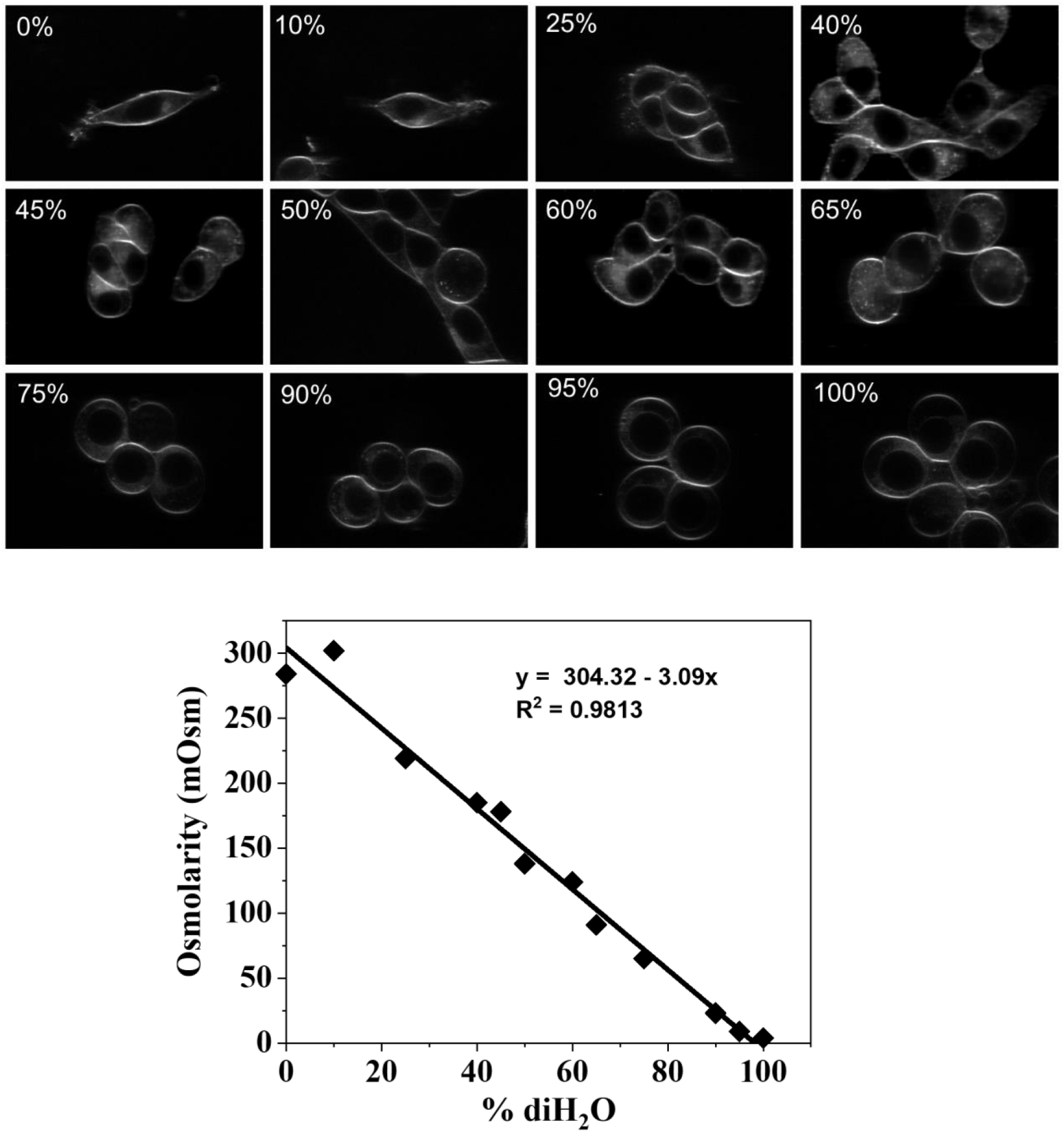
(A) Integrated fluorescence images of cells labeled with Laurdan, at different media dilutions with diH_2_O. (B) Osmolarity of the solutions as a function of diH_2_O content.

**Figure 3:**
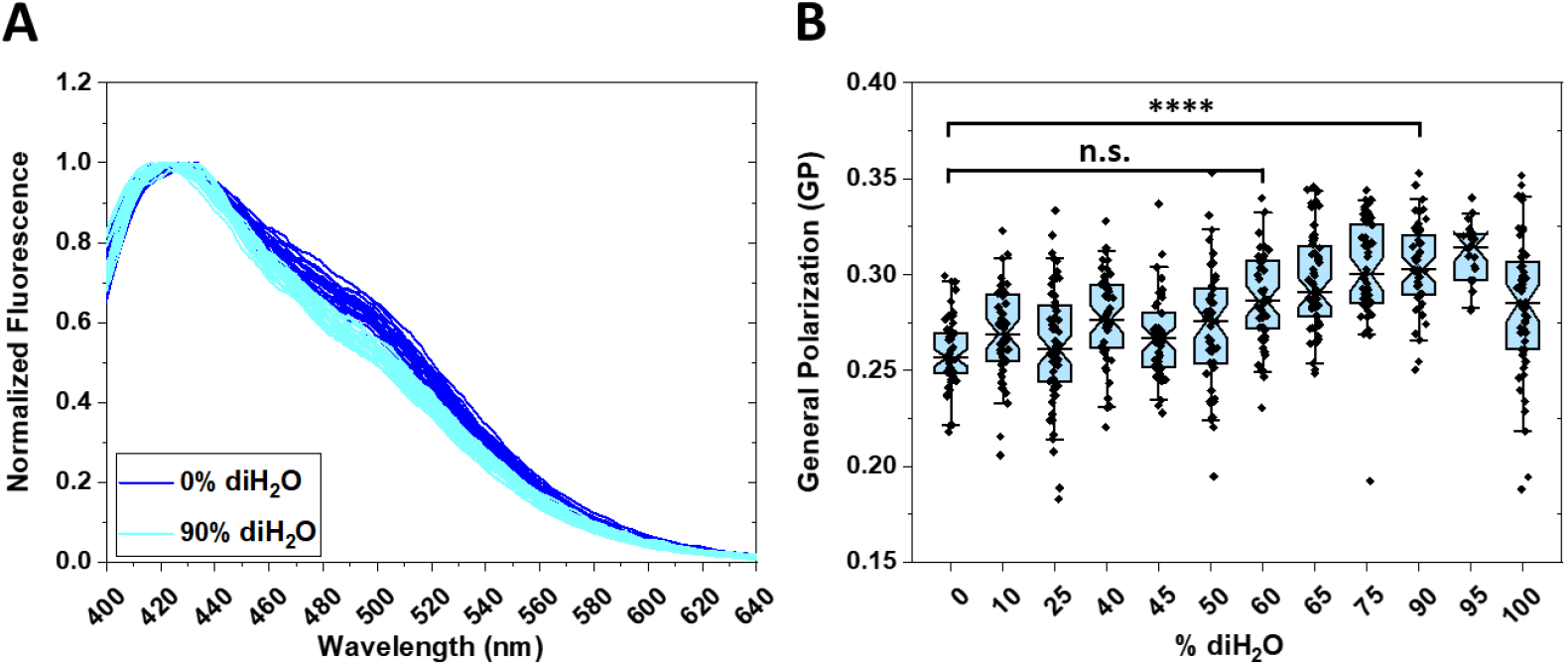
(A) Emission spectra of Laurdan in CHO cells at 0% and 90% media dilution. (B) Laurdan GP in CHO cells as a function of diH20 media dilution. GP was calculated using equation (1). The notched box line represents the mean; the top and bottom represent the 75^th^ and 25^th^ percentiles, respectively, and the bars represent the standard deviations.

Our finding that the application of osmotic stress increases GP, while the opposite was observed in lipid bilayers, was surprising. To gain confidence in our results, we performed control experiments where we increased the temperature without swelling the cells. The Laurdan spectra at room temperature (RT), 37°C, and 42°C are shown in **Figure 4A** and reveal that the amplitude of the 490nm peak increases with temperature. The calculated GP is presented in **Figure 4B** and decreases when the temperature increases. These results are consistent with the literature, as it is accepted that the increase in temperature increases its water penetration as it disorders the membrane (48,49). Thus, we find that the temperature and the application of osmotic stress have opposite effects. Furthermore, a decrease in GP was observed when the temperature was increased even in cells under reversible osmotic stress (**Figure 4C, 4D**).

**Figure 4:**
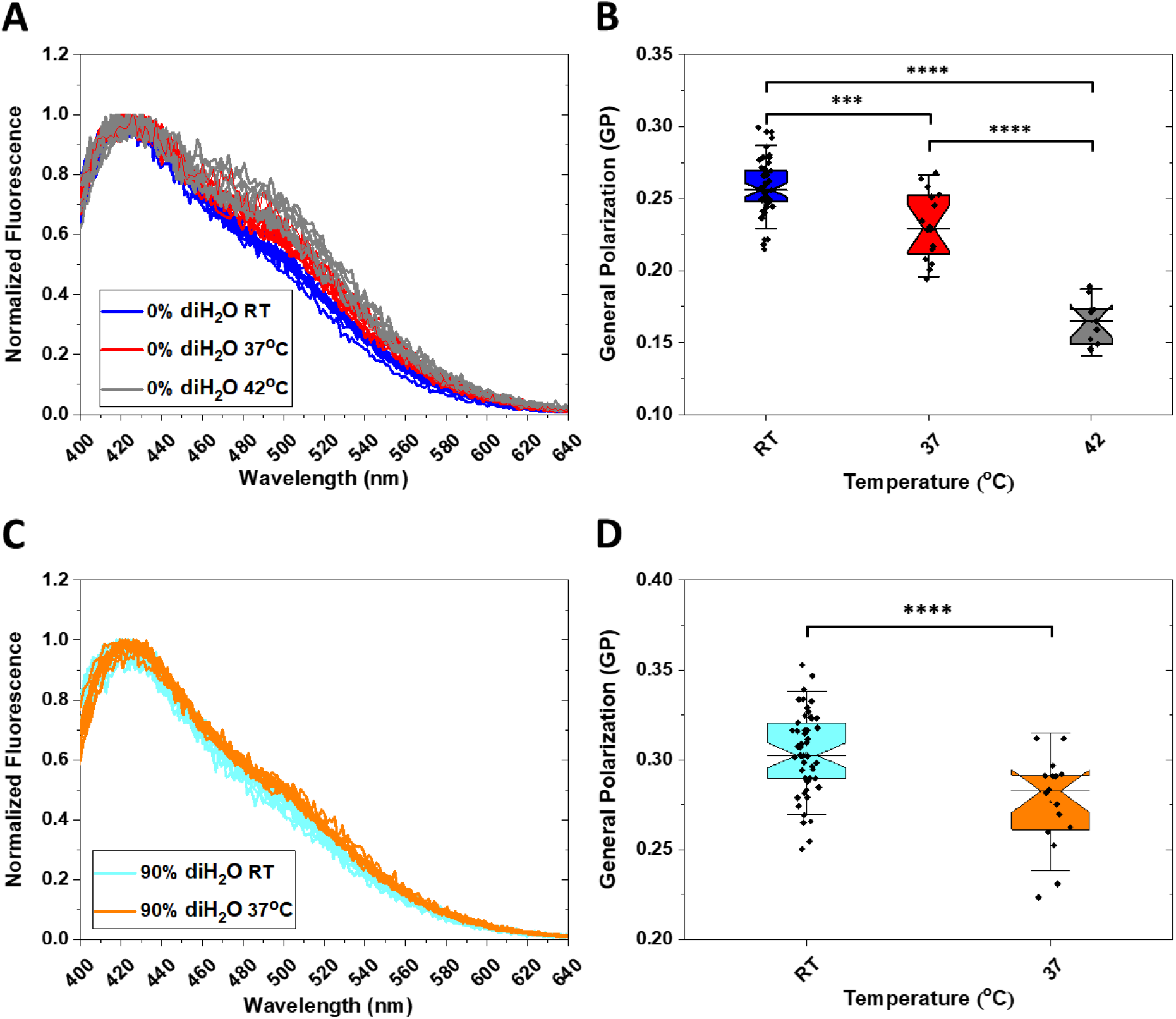
(A) Emission spectra of Laurdan in CHO cells in normal media, as a function of temperature; RT in blue, 37°C in red, and 42°C in dark grey. (B) Laurdan GP as a function of temperature in CHO cells. (C) Emission spectra in CHO cells at 90% media dilution, as a function of temperature; RT in cyan, 37° in orange. (D) Laurdan GP as a function of temperature at 90% media dilution. The notched box line represents the mean; the top and bottom represent the 75^th^ and 25^th^ percentiles, respectively, and the bars represent the standard deviations.

The application of osmotic stress has been linked to the flattening of the caveolae (8). We therefore hypothesized that our observations are intimately linked with caveolae, and thus GP would not increase if there were no caveolae. To test this hypothesis, we used Mouse Embryonic Fibroblasts (3T3 MEFs) that do not express caveolin1 and thus do not have caveolae (3T3 MEFs (-/-)). As a control, we also used the normal 3T3 MEF (+/+). Typical Laurdan spectra for the two cell lines are shown in **Figure 5A** and **5B**, for the cases of 0% and 90% media dilution with diH_2_O. We found that the GP measured for 3T3 MEFs (+/+) is increased upon the application of reversible osmotic stress, similar to the results observed in CHO cells. On the other hand, the GP for 3T3 MEF (-/-) was the same in the presence and absence of osmotic stress. Furthermore, the GP values measured in 3T3 MEF (-/-) cells lacking caveolae were different from the GP value for 3T3 MEF (+/+) cells at 0% diH_2_O, but indistinguishable from the 3T3 MEF (+/+), 0% diH_2_O GP value. These results show that the cell membrane lacking caveolae is similar to normal cell membranes under reversible osmotic stress but has higher GP values when compared to normal cell membranes in regular cell media. These data are thus consistent with our hypothesis that the GF increase is a consequence of the flattening of the caveolae or the “un-wrinkling” of the membrane.

**Figure 5:**
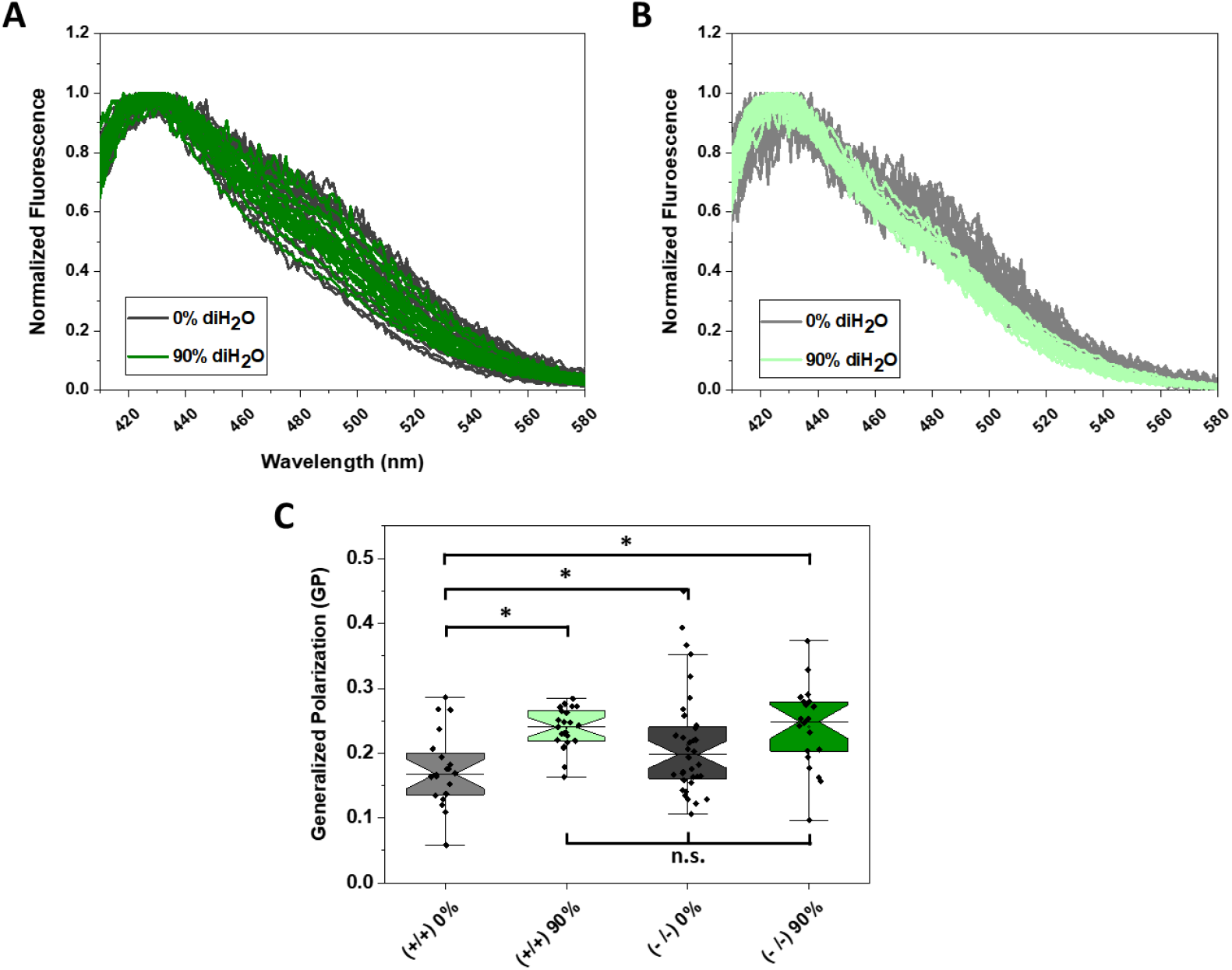
(A) Emission spectra of Laurdan in MEF (-/-) cells at 0% media dilution (dark grey) and 90% media dilution (dark green). (B) Emission spectra of Laurdan in MEF (+/+) cells at 0% media dilution (light grey) and 90% media dilution (mint green). (C) Comparison of Laurdan’s GP for MEF (+/+) and (-/-) cells at 0% media dilution and 90% media dilution. The notched box line represents the mean; the top and bottom represent the 75^th^ and 25^th^ percentiles, respectively, and the bars represent the standard deviations.

A cell under reversible osmotic stress has a topology that resembles a giant plasma membrane-derived vesicle (50). Such vesicles are produced by cells in response to the so-called vesiculation buffers and are increasingly being used in biophysical research (51-53). We used Laurdan as a probe to compare the membrane environment of cells under reversible osmotic stress and vesicles produced with the widely used formaldehyde method. The results are shown in **Figure 6**. They demonstrate that there are no statistically significant differences between GP values of swollen cells and plasma membrane vesicles, suggesting that they have a similar membrane environment.

**Figure 6:**
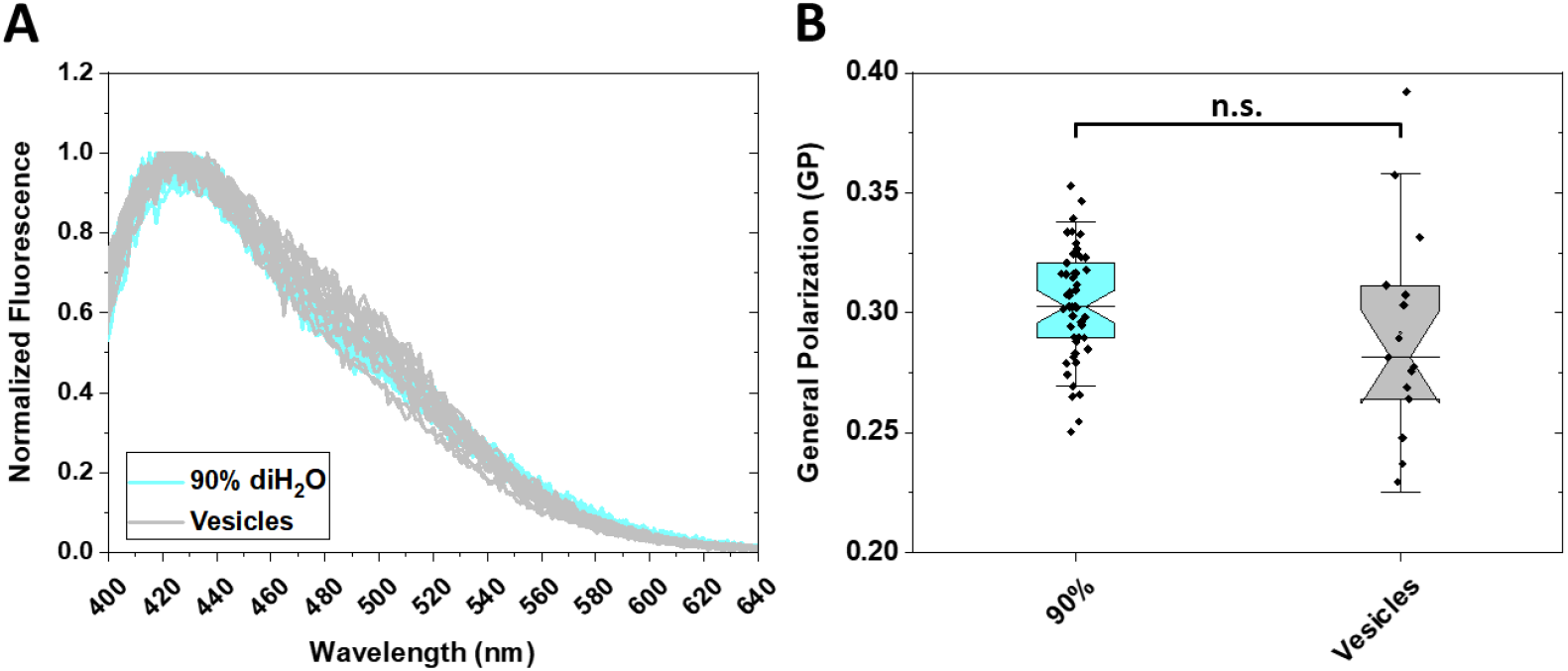
(A) Emission spectra of Laurdan in CHO cells at 90% media dilution (cyan) and in plasma membrane-derived vesicles (light grey). (B) Comparison of Laurdan GP for CHO cells at 90% media dilution and membrane-derived vesicles. The notched box line represents the mean; the top and bottom represent the 75^th^ and 25^th^ percentiles, respectively, and the bars represent the standard deviations.

## Discussion

The main finding in this paper is that Laurdan’s GP, measured in the plasma membrane, increases upon the application of reversible osmotic stress. This increase in GP occurs only at high osmotic stress once the cell media is diluted by 50% or more with diH_2_O. The effect is opposite to what is observed upon increase in temperature. Furthermore, the effect of osmotic stress is different in the plasma membranes of live cells and in lipid vesicles which are often used as models of the plasma membrane. The GP has been found to decrease in lipid vesicles in response to osmotic stress (44), and this decrease was interpreted as an indication of bilayer stretching, which leads to an increase in water penetration (44).

In the laboratory, researchers use reversible osmotic stress to flatten the caveolae and thus “un-wrinkle” the plasma membranes in a reversible manner, such that the membrane topology is known in biophysical experiments (54,55). Indeed, in fluorescence microscopy experiments, the application of osmotic stress results in spherical plasma membrane topology, where the membrane is perpendicular to the imaging plane. In this case, the two-dimensional concentration of fluorophores in the plasma membrane can be calculated from the effective 3D concentrations through a multiplication by the pixel length. The application of reversible osmotic stress to cells is non-lethal and is fully reversible (8,50,55,56). Indeed, we have shown that once the swelling medium is replaced with starvation medium, and the cells are placed back in the incubator, they recover fully, consistent with other reports of complete reversibility (34). Furthermore, the FRET efficiencies measured for membrane proteins are not affected by the application of reversible osmotic stress (56), suggesting that the interactions are very similar in unperturbed cells and in cells under reversible osmotic stress. However, a concern may arise that the lipid membrane is stretched due to the application of osmotic stress. This concern motivated the current study. In our cellular study, a decrease in GP was observed only at the highest osmotic stress when the diH_2_O content was increased above 95%. Under these conditions, the plasma membranes often ruptured during the imaging session, indicating that pressures leading to membrane stretching were not well tolerated by cells. However, we found that cells could easily withstand media dilution up to 90% with no indications for membrane stretching.

How can we explain the GP increase upon the application of osmotic stress? One possibility is that it is a direct consequence of hydration changes in the caveolae. In particular, the necks of the caveolae are sites of very high curvatures, imposed by the organizing proteins. It is possible that these regions are highly strained and hydrated, and the lipids are highly disordered. It could be speculated that the application of osmotic stress leads to highly localized changes in the necks of the caveolae that expel water molecules. Since these areas are much smaller than the diffraction limit of light, we likely observe an average increase in the GP in the membrane. Another possibility is that the GP in our experiments reports mainly on changes in lipid mobility in the membrane upon the application of osmotic stress, with an increase in GP indicating decreased mobility. It is known that the lipid composition of caveolae is enriched in cholesterol and sphingolipids, and thus caveolae are regions of low lipid mobility (57-59). Upon caveolae disassembly, the prevalence of cholesterol and sphingolipids in these regions should decrease. However, this effect cannot explain our observations as it would lead to an increase in GP, i.e., the opposite effect of what is observed. It is possible, however, that the dispersal of cholesterol and sphingolipids from the caveolae into the membrane has a marked effect on lipid mobility which we measure here. Further, it should be noted that cavoelae play a role in signaling (57), and thus downsream signalimg effects may contribute to the observations reported here. Future studies are needed to fully understand all cellular events that contribute to the observed increase in GP.

GP is a phenomenological parameter, often used in membrane biophysics since steady-state Laurdan fluorescence is straightforward to measure. While the interpretation of GP may pose a challenge, the fact remains that the change in GP upon the application of osmotic stress is exactly the opposite in lipid vesicles and in cells. Thus, the behavior observed here in cells could not be predicted based on measurements in model systems.

Lipid bilayers have been an extraordinarily useful model system for understanding processes in cellular membranes. This study reminds us that measurements in this simple system cannot be expected to always mirror the response of the plasma membrane. Likewise, plasma membrane-derived vesicles are increasingly being used in biophysical research (51,60). Our results suggest that they represent a membrane environment with similar physical-chemical properties as the plasma membrane in cells under reversible osmotic stress, as well as in cells with no caveolae. It is also interesting that cells can easily withstand very high media dilution with no indications for membrane stretching. This work contributes to our understanding of membrane models and the limitations they impose, as we seek to understand processes in native plasma membranes using biophysical tools.

## Acknowledgement

Supported by NSF MCB 1712740. We thank Dr. Anne K. Kenworthy for reading an early version of the manuscript, and for suggesting the 3T3 MEF (-/-) line as control. We thank Brandon Tenney, Maryann Ness, and Mark Marzinke from the Johns Hopkins Hospital Core Lab facility for the osmotic measurements.

## Notes

### Competing Interest Statement

The authors have declared no competing interest.

